# Contemporary risk of extinction in an extreme environment

**DOI:** 10.1101/2020.09.15.298919

**Authors:** Simone Vincenzi

## Abstract

The increased frequency and intensity of extreme events are recognized among the most worrisome aspects of climate change. However, despite increased attention from scientists and conservationists, developing and testing general theories and hypotheses on the effects of extreme events on natural populations remains intrinsically challenging.

Using numerical simulations with general—but realistic for moderately fast-leaving species—parameter values, I tested some of the hypotheses on risk of extinction and population and genetic dynamics in an environment in which both climate (e.g., temperature, rainfall) and point (e.g., fires, floods) extremes occur. In the simulations, a quantitative trait is selected for by a climate variable, but point extremes cause trait-independent massive mortalities.

I found additive effects between age at first reproduction and fecundity on risk of extinction. The extent of population bottlenecks (operationally, the number of years in which a population was at low numbers) was a good predictor of allelic richness for the quantitative trait selected for by the climate. Simple models including basic demographic and vital rates information of the species, along with climate/environmental measures, provided excellent predictions of contemporary risk of population extinction. Mean and minimum population size measured in a 10-year “observation window” were largely the most important predictors of risk of population extinction in the following 10-year “prediction window”.

## 1 Introduction

Extreme events are now recognized among the most worrisome aspects of climate change (IPCC 2007, 2012, Jentsch et al. 2007, The One Earth Editorial Team 2020). They may be defined in terms of extreme values of a continuous climate variable (e.g., daily or mean summer temperature, rainfall) on the basis of the available climate and weather record (Gutschick and BassiriRad 2010), or in the form of a “point” (or discrete) perturbation, such as a hurricane or a storm; the latter category also includes environmental hazards such as fires and floods. Following Vincenzi (2014), throughout this work I will use the term climate extreme for extreme values of a continuous climate or environmental variable and point extremes for discrete extreme perturbations.

The genetic and demographic determinants of both adaptation to a more extreme environment and contemporary risk of extinction are receiving increasing interest from scientists and conservationists. Despite the increased research and management focus on extremes, developing and testing general theories and hypotheses on the effects of extreme events remains more challenging than for other aspects of climate and environmental change (van de Pol et al. 2017). Those challenges are likely to be intrinsic and unavoidable.

First, since the effects of extreme events are largely context-specific, developing an overarching causal and predictive framework might be overly ambitious (Bailey and van de Pol 2016). For instance, the emergence of adaptations may depend on both the life histories of the species and the recurrence interval, intensity, and nature of extreme events. Besides, the demographic and genetic effects of climate and point extremes (e.g., population crashes, loss of genetic diversity, inbreeding and maladaptation, changes in population age and size structure, disruption in the expression of quantitative traits) (Bryant and Meffert 1995, Kirkpatrick and Jarne 2000) are often the result of chance and are thus not easily predictable or generalizable across species or habitats. Second, climate and point extremes that result in strong demographic and genetic responses are often rare events, whose occurrence may be also difficult to predict. When available, pre-disturbance empirical data is likely to have been collected by chance, and the studies on the effects of extreme events prone to be mostly opportunistic and anecdotal instead of planned and comprehensive.

Despite the intrinsic difficulties of developing and testing general theories on the effects of climate and point extremes on natural populations, there are a few theory-based predictions on the demographic and genetic consequences of extreme events that may be quite general across species and habitats, although with varying degrees of modeling and empirical validation.

The first prediction is that by reducing population size, extreme events directly increase the risk of extinction of the affected populations (Willi et al. 2006, Frankham et al. 2014); mathematical modeling, empirical observations, and common sense all suggest that smaller populations are more likely to go extinct than larger populations. After the occurrence of extreme events, in some cases the extinction of the focal population is inevitable for numerical reasons (no individuals capable of reproducing survived) unless there is immigration from neighboring—either unaffected or less affected—populations (i.e., “demographic rescue”, Brown and Kodric-Brown 1977, Carlson et al. 2014).

Second, when the change in the environment is sudden, but not causing mass mortalities, the evolution of fitness-determining traits might occur fast enough to stop population decline and allow population recovery before extinction (“evolutionary rescue”, Bell and Gonzalez 2009). However, population bottlenecks such as those caused by extreme events, especially when repeated over time, are predicted to decrease additive genetic variance and allelic diversity in the affected populations (Bouzat 2010). As adaptive potential tends to increase with genetic variability and genetic drift may overwhelm selection in small populations, the effects of extreme events on genetic variability are predicted to increase both the short- and long-term risks of population extinction, with smaller chances of evolutionary rescue (Falconer and Mackay 1996, Willi and Hoffmann 2009).

Third, species may exhibit various adaptations to extreme events, some more predictable and general than others. For instance, extreme, but predictable variations in the flow regime of streams can select for fish life histories, such as spawning and emergence time, that are synchronized either to avoid or exploit the direct (e.g., stronger or weaker currents) and indirect (e.g., changes in food webs) effects of extreme flows (Lytle and Poff 2004). On the contrary, unpredictable flow events such as flash floods may have low direct selective consequences for the affected populations, even though they might induce massive mortalities (Lytle 2000). In these cases, natural selection after the extreme event is predicted to favor individuals with a high capacity for increase in population size *(r* selection, Reznick et al. 2002). Intuitively, when after an extreme event the population is reduced to a few individuals, faster reproduction may be more critically needed than high fecundity, since the latter depends on being able to reproduce. However, the interaction effects between age at first reproduction and fecundity on risk of extinction in an extreme environment have been rarely investigated.

In this work, I test using a simulation approach some of the hypotheses on risk of extinction and population and genetic dynamics in an environment in which both climate and point extremes occur. In previous work, Vincenzi (2014) found that the survival chances of a population were found to decrease with increasing strength of selection, as well as with increasing climate trend (e.g., increasing or decreasing *n*-year moving average temperature, rainfall) and variability. They also found that the interactions among climate trend (e.g., increase over years of average summer temperatures), climate variability and probability of point extremes (e.g., fires) had negligible effects on risk of extinction, time to extinction, and distribution of a quantitative trait selected for by climate after accounting for their independent effects.

The present work focuses more on prediction than inference; prediction of future observables has long been included as an aspect of biological and ecological studies, but as a methodological approach it has been much less prominent than either description or statistical and causal inference. The work is also not purely theoretical in scope, but is motivated (and model parameter values inspired) by the study of the population and genetic dynamics, and risk of extinction of marble trout *Salmo marmoratus* living in Slovenian streams (Vincenzi et al. 2016). Marble trout populations are affected by flash floods in autumn and droughts in summer, and the climate threat to the persistence of marble trout is likely to worsen with climate change; for instance, the increase in intra-annual variability in rainfall is predicted to increase the frequency of flash floods (Janža 2013). However, since there is a need for more general investigations of risk of population extinction and population and genetic dynamics in presence of climate and point extremes, I developed a more widely applicable modeling and conceptual framework.

I modeled an idealized situation of additive genes, a closed population of moderate size with random mating, and variable across simulations—but fixed within simulations—age at first reproduction and expected number of offspring per mating pair, climate trend and variability (i.e., the parameters of the distribution of climate variable that is selecting for a quantitative trait), selection strength, and frequency and severity of point extremes. I assume that point extremes cause massive mortalities, but no trait other than good fortune increases the survival chances of an individual when point extremes occur.

First, I tested whether in an extreme environment the extent of population bottlenecks, which I operationally defined as the total number of years with “depressed” population size (e.g., ~ 1/3 to ~ 1/10 of the habitat carrying capacity), could help predict allelic richness at the end of simulation time for the quantitative trait selected for by the climate variable. Second, I tested whether quantitative trait adaptation was correlated with the increased frequency of theoretically advantageous alleles (i.e., their allelic values are in the same direction of the change in the environment) and whether the genetic dynamics of the populations could be predicted using information on the climate and on the population life histories. Third, I tested for interaction effects between age at reproduction and fecundity on risk of population extinction. Finally, I tested whether it is possible to predict the extinction or survival of a population in a 10-year “prediction window” (i.e., contemporary extinction) when measuring or observing some of the environmental characteristics of the habitat (e.g., occurrence of extreme events) and of the population (e.g., population size, age at reproduction) during a 10-year “observation window” that immediately precedes the “prediction window”.

## 2 Material and methods

The model I use in this work is an extension of the model developed in Vincenzi (2014). The choice of parameter values for the present work was informed by the available literature on extreme events, population monitoring, and species life histories, and by the results of Vincenzi (2014).

### 2.1 Overview of the model

I consider a population of monoecious individuals living in a habitat whose population ceiling is *K* (Mangel and Tier 1993). The population is geographically isolated, with neither immigration nor emigration from or to other populations. A single quantitative trait *a* corresponding to its breeding value for a phenotypic trait *z* characterizes the individuals. The population has discrete overlapping generations with *N*(*t*) total individuals, where *t* is time in years. The environment is described by an optimum phenotype *Θ*(*t)* that changes over time as a result from variations in a climate driver such as rainfall or average summer temperature, which selects for the trait *z*. The distance between the optimum phenotype *Θ(t)* and the trait *z*_i_, (*Θ(t)* - *z*_i_) of the individual *i* defines the maladaptation of the individual *i* with respect to the optimum phenotype (or, alternatively, it defines the “extremeness” of the climate event for the individual). Point extreme events such as floods or fires cause non-selective high mortality in the population, i.e. every phenotype has the same chances of surviving the event.

### 2.2 Optimum phenotype

The expected optimum phenotype *μΘ*(*t*) moves at a constant rate *β_μ,Θ_* over time (i.e., trend), fluctuating randomly around its expected value *μ_Θ_(t)*. The optimum phenotype *Θ(t)* is randomly drawn at each time step *t* from a normal distribution ***Θ***(*t*) ~ *N* (*μ_Θ_*(*t*),*σ_Θ_*(*t*)). It is equivalent to consider *Θ(t)* as both the optimum phenotype and the value of a continuous climate variable (e.g., mean summer temperature or yearly rainfall), and I will use the two terms interchangeably throughout this work.

Mean and variance of the climate variable at time *t* are thus:

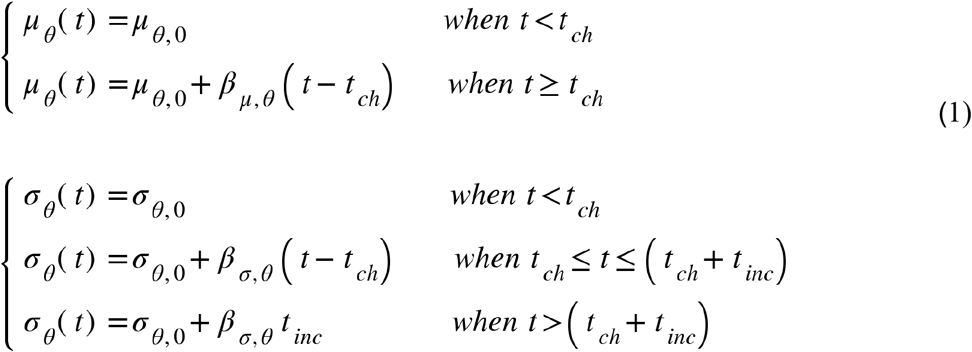

where *t*_ch_ is the time at which there is a change (*ch*) in the climate. Eq. (1) indicates that the directional climate trend steadily increases through time after *t*_ch_ years and that the increase in variability starts after *t*_ch_ years, but stops after *t*_ch_+*t*_inc_ years. An alternative formulation to Eq. (1) would allow variability to more slowly build up over time; however, in the case of variability slowly increasing over longer times, the effects of higher climate variability would affect the individuals only towards the end of simulation time. As informed by the goals of this work, I preferred to let individuals trying to survive a higher climate variability (which is among the least investigated aspects of climate and environmental change) for a longer time period.

With the model formulation of Eq. (1), both the mean and variance of the distribution of the climate variable change over time so as to make the occurrence of events more likely after climate change (i.e., *t*>*t*_ch_) than before, since after climate change, the realized (i.e., random draw from the statistical distribution of climate) climate is increasingly likely to be in the region of extremes (say, in the right 5% or 2.5% of the Gaussian distribution of climate) of the statistical distribution of climate before climate change.

Point extreme events *E* leading to trait-independent high mortalities occur with annual probability p(*E*_b_) when *t*<*t*_ch_ (i.e. b - before climate change) and p(*E*_a_) when *t*>*t*_ch_.

### 2.3 Quantitative trait and survival

I model the phenotype *z* of an individual *i, z*_i_, as the sum of its genotypic value *a*_i_ and a statistically-independent random environmental effect *e*_i_ drawn from 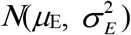:

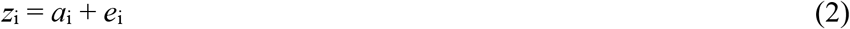

The narrow sense heritability 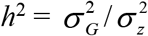 is the proportion of the phenotypic variance 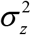 present in the population that is explained by the additive genetic variance 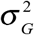 (i.e. the variance of ***a*** in the population).

For an individual *i*, the genetic value *a*_i_ is determined by *n*_1_ freely recombining diploid loci, with additive allelic effects within- and among-loci, that is 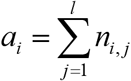, where *n_i, j_* is the sum of the allelic values at locus *j*. For computational reasons, I chose *n*_1_ = 10. Allelic values are randomly drawn from a Gaussian distribution with mean of 0 and variance equal to 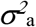. For simplicity, I did not model either dominance or epistatic variation or other complicating factors such as genotype-environment interaction and linkages. Likewise, I did not model mutation, since previous work has shown that mutation does not appear to have any effect short-term on extinction risk and the evolution of traits on contemporary temporal scales (Vincenzi 2014).

Stabilizing selection is modeled with a Gaussian function (Bürger and Lynch 1995, Zhang 2012), with fitness *W* (Endler 1986) for an individual with phenotypic trait *z*_i_ equal to:

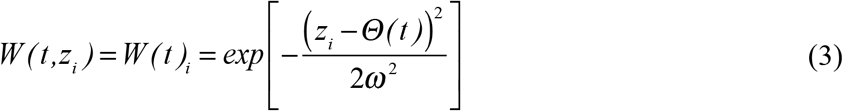

and equivalent in this model to the annual survival probability of individual *i*. The curvature of the fitness function near its optimum increases with decreasing *ω*^2^; it follows that that the smaller *ω*^2^, the stronger is selection. Stabilizing selection is usually measured by the standardized quadratic selection gradient *γ*, which is defined as the regression of fitness *W* on the squared deviation of trait value from the mean (Lynch and Walsh 1998). An optimum phenotype in the tails of the distribution is likely to cause a large drop in population size and can be considered an extreme climate event (Fig. 1).

**Figure 1.**
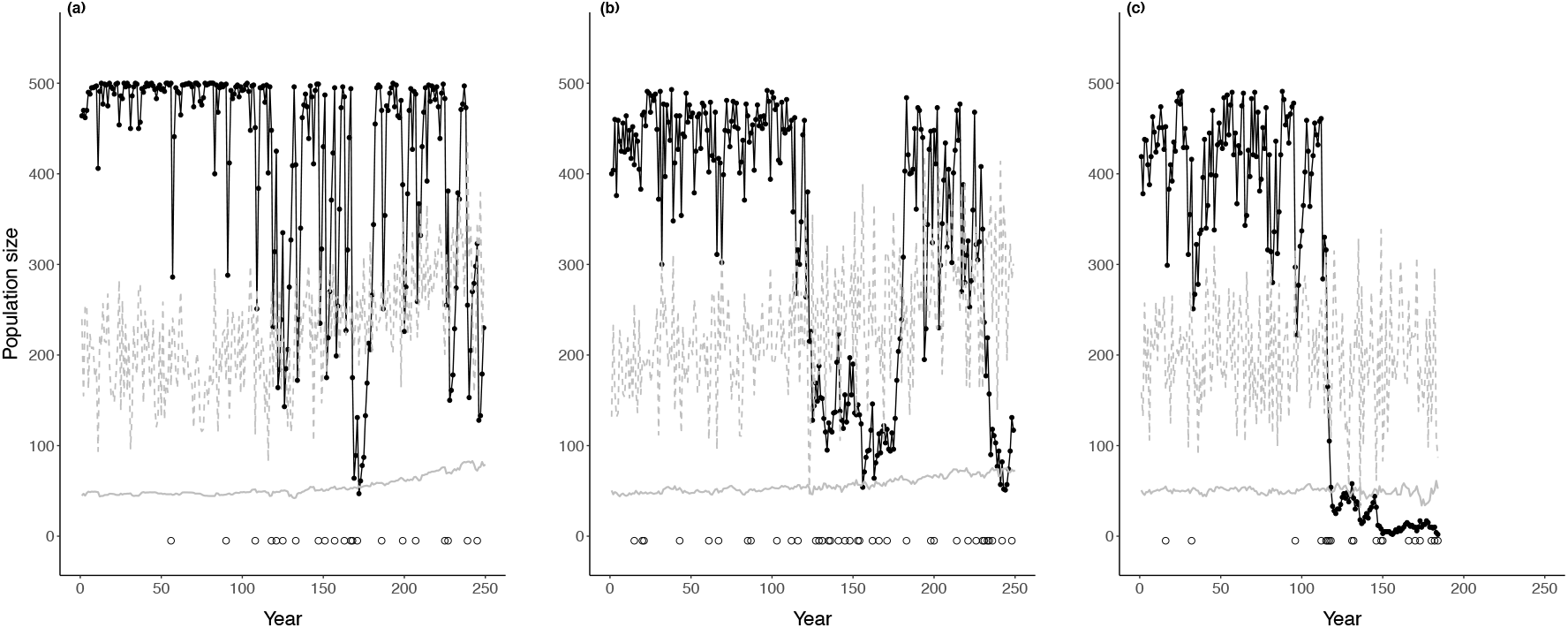
Examples of simulation replicates. The black solid line and black points represent population size, circles are point extremes, the gray dashed line is the value of the optimum phenotype *Θ*, and the solid gray line is the mean phenotype 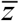 in the population. Both *Θ* and 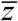 are re-scaled for graphical purposes. Parameters for the simulation in panel (a): *β_σ,Θ_* = 0, *β_μ,Θ_* = 15 10^-2^, p(*E*_a_) = 10 10^-2^, *m*_E_ = 0.7, *a*_f_ = 4, *λ*0 = 2, *s* = 11 10^-2^. (b): *β_σ,Θ_* = 15 10^-2^, *β_μ,Θ_* = 15 10^-2^, p(*E*_a_) = 15 10^-2^, *m*_E_ = 0.3, *a*_f_ = 3, *λ*_0_ = 2, s = 8 10^-2^. (c): *β_σ,Θ_* = 15 10^-2^, *β_μ,Θ_* = 0, p(*E*_a_) = 15 10^-2^, *m*_E_ = 0.5, *a*_f_ = 3, *λ*_0_ = 1, *s* = 8 10^-2^. In the simulation in panel (c), the population went extinct at year 184. Symbols are as in Table 1.

The median *γ* = −0.1 for stabilizing selection found by Kingsolver et al. (2001) corresponds to a value of 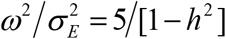, where 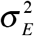 is the variance of the environmental component of the phenotype defined in Eq. (2), when stabilizing selection is modeled using a Gaussian fitness function.

Eq. (3) can be written:

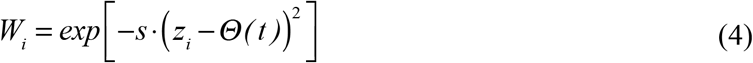

where 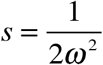. With *γ* = −0.1, 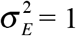, and *h*^2^ = 0.2, the strength of selection *s* is about 0.08.

I assumed that both strength of selection *s* and environmental variance 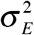 remain constant through time. When a point extreme occurs, the probability of yearly survival of individuals *i* is *W_i_(*1−*m_E_I__)*, where *m_E_I__* is mortality caused by the point extreme event.

### 2.4 Simulations

As this study focuses on the more immediate effects of climate change, the simulations last 250 years. Parents mate at time *t*-1, offspring are born at time *t* and become of age 1 at *t*+1. The sequence of operations is mortality of adults, mating and reproduction, mutation, mortality of offspring. At the start of each simulation, for each individual a value of *a* and *e* (Eq. 2) is randomly drawn from their initial distribution. A population is considered extinct if at any time during the simulation there are fewer than 2 individuals in the population. Parents form mating pairs starting at age *a*_f_ and produce a number of offspring randomly drawn from a Poisson distribution with intensity *λ*_o_. Offspring receive for the same locus one allele from each parent.

#### 2.4.1 Parameter values

With numerical simulations, it is inevitable to face a trade-off between specificity and generalizability of the modeled processes and of the simulation results. I reduced parameter space by fixing *K* = 500 individuals, 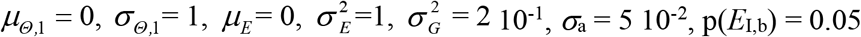, and *t_inc_* = 25. For the other parameters, I chose range of values that are both realistic for natural populations with moderately fast life histories, like salmonids (e.g., age distribution skewed toward individuals younger than 10 years old, sexual maturity reached when individuals are between 1 and 4 years old), and instrumental for the main goal of the study, e.g., testing hypotheses on the effects of extreme events on population and genetic dynamics and on risk of extinction.

I performed simulations with selection strength *s* equal to either 8 10^-2^ (average selection strength) or 1.1 10^-2^ (moderately strong selection). For the rate of increase in the mean of the climate variable, I used *β*_*μ*,Θ_ = 0 (base scenario) and 1.5 10^-2^. I used rates of the increase in the standard deviation of the climate variable *β*_*ρ*,Θ_ from 0 (base scenario) to 1.5 10^-2^. According to Bürger and Lynch (1995), when the standard deviation of the distribution of the optimum *σ_θ_* reaches the same order of magnitude as the width *ω* of the fitness function, the population is at risk of going suddenly extinct, with little role played by genetics. Therefore, I chose values of *β*_*ρ*,Θ_ that strongly increase the probability of climate extremes, but did not inevitably make the population go extinct.

I used frequency of point extreme events p(*E*_a_) of either 5 10^-2^ (no variation before and after climate change, corresponding to a recurrence interval of 20 years), 10 10^-2^ (i.e., recurrence interval is 10 years) or 15 10^-2^ (Table 1). I used moderate mortalities caused by point extremes (simulations with *m_E_* equal to either 0.3, 0.5, 0.7) and moderate p(*E*_a_), since with higher mortality induced by point extremes and higher probability of their occurrence the system will be largely driven by the point extremes, with no or little role of genetics and demography in determining population dynamics and risk of extinction.

**Table 1.** Values of parameters of the model of population and genetic dynamics.

| Parameters | Values | Description |
| --- | --- | --- |
| $K$ | 500 | Population ceiling |
| $\lambda_o$ | 1.0, 1.5, 2.0, 2.5 | Intensity of the Poisson distribution of yearly number of offspring per mating pair |
| $a_f$ | 1, 2, 3, ,4 | Age at first reproduction |
| $t_{ch}$ | 100 | Years since the start of the simulation before climate change |
| $t_{inc}$ | 25 | Time of increase of variability (variance of the normal distribution of the climate variable) after climate change |
| $n_l$ | 10 | Number of diploid loci |
| $\sigma_A^2$ | $6.25 \cdot 10^{-3}$ | Additive genetic variance per locus at the start of simulation |
| $\sigma_a^2$ | $2.5 \cdot 10^3$ | Variance of the normal distribution of allelic values |
| $\sigma_G^2$ | 0.2 | Additive genetic variance of the quantitative trait at the start of simulation |
| $\mu_E$ | 0 | Mean environmental effect |
| $\sigma_E^2$ | 1 | Environmental variance |
| $m_E$ | 0.3, 0.5, 07 | Mortality caused by the point extreme event |
| $s$ | 8, 11 $10^{-2}$ | Strength of selection |
| $p(E_b)$ | $5 \cdot 10^{-2}$ | Probability of occurrence of point extreme events before climate change |
| $p(E_a)$ | 5, 10, 15 $10^{-2}$ | Probability of occurrence of point extreme events after climate change |
| $\mu_{\theta,0}$ | 0 | Mean of the normal distribution of the phenotypic optimum from year 1 to $t_{ch}$ |
| $\sigma_{\theta,0}$ | 1 | Standard deviation of the normal distribution of the phenotypic optimum from year 1 to $t_{ch}$ |
| $\beta_{\mu,\theta}$ | 0, 1.5 $10^{-2}$ | Annual increase (directional trend) of the mean of the normal distribution of the climate variable from year $t_{ch}$ to the end of simulation |
| $\beta_{\sigma,\theta}$ | 0, 1, 1.5 $10^{-2}$ | Annual increase of the standard deviation of the normal distribution of the climate variable from year $t_{ch}$ to $t_{ch} + t_{inc}$ |

For the Poisson distribution of the yearly number of offspring per mating pair, I used *λ*_o_ equal to either 1.0, 1.5, 2.0 or 2.5 and age at first reproduction from 1 to 4 years old with a step of 1. Parameter values are reported in Table 1.

#### 2.4.2 Initialization

To reach mutation-selection-drift balance, I first let the population evolve for *t*_ch_ years in an environment in which mean and variance of the distribution of the optimal phenotype ***Θ*** are constant. In preliminary simulations it was found that after *t*_ch_ ~ 100 years both phenotypic mean and variance did not noticeably change. Then, the mean of ***Θ*** increases for 150 years and the variance of ***Θ*** for 25 years, which was then kept constant up to the end of simulation time.

I started every simulation replicate with 500 individuals. I modeled 10 alleles present in the population for each locus, which value was randomly drawn from a normal distribution 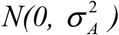. Since I set 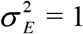 and 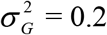, the narrow sense heritability *h*^2^ was around 0.2 at *t* = 1, close to what commonly observed for life-history traits (Lynch and Walsh 1998) and consistent with the Gaussian allelic approximation including only quasi-neutral and adaptive mutations, for which 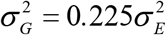 (Lande 1995).

#### 2.4.3 Characterization of simulations

At the single-replicate level, to characterize the behavior of the simulated populations I tracked or recorded (among other results): (*a*) whether the population was extinct or still persisting at the end of the simulation time (0 for persistence and 1 for extinction, in the latter case I also recorded the year of extinction). At each time *t*, I then recorded: (*b*) the distribution of the trait *z* in the population and individual maladaptation; (*c*) population size *N* after mortality of adults; (*d*) total number of alleles and allelic frequencies (the latter every 25 years).

For an ensemble of realizations (10 replicates for a fixed set of parameters) I also computed the frequency of population extinction as the number of replicates in which the population went below two individuals during simulation time.

### 2.5 Statistical analysis

I used simulation results as pseudo-empirical data and analyzed them with standard statistical and machine learning models. The main focus of the statistical analyses and modeling was more on prediction than on inference or traditional *p*-value hypothesis-testing.

I estimated parameters of Generalized Additive Models (GAMs, Wood 2006), Generalized Linear Models (or Ordinary Least-Square regression models, McCullagh and Nelder 1989), and Random Forests (Breiman 2001a) using as response variable either (*i*) the total number of alleles (overall allelic richness) at the end of simulation time for the populations that persisted, (*ii*) the difference in mean allelic frequency of the top 10% and bottom 10% (according to their allelic value, top 10% were the alleles with the bigger allelic value and *vice versa* for the bottom 10%) of alleles. For (*i*), in particular, I tested whether the number of consecutive years (*n_low*, the only variable in the GAM models for which I hypothesized a non-linear effect on the total number of alleles) under a low population size threshold (*tr_low*) contributed to predicting the number of alleles at the end of simulation time, in addition to climate variables and vital rates such as expected intensity and frequency of extreme events, the variance of the optimum, expected yearly number of offspring per mating pair and age at first reproduction. I calculated *tr_low* using either 150, 100, or 50 individuals as the threshold (results were consistent when using either 150,100, or 50 individuals). For (*ii*), when the climate is changing (i.e., *β_μ,Θ_* > 0), alleles with positive allelic values are expected to be more adaptive than those with negative (or positive, but smaller) allelic values. I tested whether a model including climate variables and vital rates trained on data from 80% of the replicates that did not go extinct could predict the difference in mean allelic frequency at year 250 (end of simulation time) of the top and bottom 10% of alleles according to their allelic values in the test data set (20% replicates that did not go extinct). I chose the top and bottom 10% of the alleles (10 alleles in the top and bottom sets), since the fate of single alleles is more likely to be affected by chance than that of a group of advantageous alleles.

Then, I investigated whether a combination of demographic and environmental factors measured or estimated in a short time window (“observation window”) can predict the risk of extinction of the population in the following years (“prediction window”). First, I set aside a balanced test data set of simulation replicates (50% that went extinct and 50% that survived)—these simulations were not used in any phase of either model development or model training. Second, since the number of simulations that went extinct was approximately one-third of the number of those that survived, I augmented the data set by replicating 3 times the simulations that went extinct and were not included in the test data set.

Third, I fitted GLMs and GAMs with binomial error distribution (i.e., logistic regression), and classification Random Forests (RFs) with population extinction (1) or persistence (0) between (*t*_ext_ - *u)* as response variable, where *t*_ext_ is either (a) the time at extinction for the replicate that went extinct, or (b) a random deviate from a uniform distribution bounded between 20 and 250 for the replicates that survived; *u* is a random deviate from a uniform distribution bounded between 1 and 10 years. This way, I am trying to model extinction or persistence not at a specific time, but in a “prediction window” of 10 years, whose beginning lays between year 10 and year 240 of simulation time. I used as candidate predictors, as measured in the 10 years (“observation window”) before the “prediction window”, minimum and mean population size *N*, the maximum value of the optimum phenotype, the expected yearly number of offspring per mating pair, age at first reproduction, and maximum and mean distance over the observation window between the mean phenotype and the optimum (i.e., maximum and mean population-level maladaptation, or maximum and mean “extremeness” of the climate).

In other words, I wanted to test whether a model including climate and population traits measured over a limited time frame could predict the extinction or persistence of the population in the years immediately following the end of the “observation window”. The choice of the length of both the observation and prediction windows was based on what it is considered a “reasonable” length of population monitoring when the goal is to detect changes in population size and estimate the risk of population extinction. White (2019) collected the time series of population size of 822 populations of vertebrate species and found that 72% of time series required at least 10 years of continuous monitoring to achieve a high level of statistical power to detect significant trends in abundance (the code associated with this work allows to use between 5 and 20 years as length of the observation and prediction windows).

For the GAMs and GLMs, I estimated the optimal cutoff given equal weight to sensitivity (the probability that the model predicts extinction when the replicate went extinct) and specificity (the probability that the model predicts persistence when the replicate persisted). Then, I tested the model by predicting population extinction and persistence on the test dataset using the computed optimal cutoff. For the classification RF models, I directly used the binary prediction (population going extinct or surviving) from the models. I used different modeling approaches because I did not explicitly model any mechanism or process that can be hypothesized to lead to extinction (i.e., models are correlative and not mechanistic), and different modeling approaches can give different insights on how contemporary extinctions are predicted (e.g., tree-based models like RFs provide a measure of variable importance for predicting the target variable (Breiman 2001a) and GAMs can model non-linear relationships between predictors and target variable using semi-parametric estimation). For the GAMs and GLMs, I centered and scaled the predictors in order to compare their importance (Schielzeth 2010). As I use realistic variable ranges representing the variability that may be observed in nature, some of the estimated parameters can be compared in terms of effects on a standardized scale. I also fitted the same models using non-standardized predictors to test for possible data leakage between training and testing data sets (results were directionally the same). I did not include interactions among predictors in the models in order to improve the interpretability of results. I visually checked residuals for violation of model assumptions.

## 3 Results

Results are fully reproducible. Data and R code are at https://github.com/simonevincenzi/Contemporary_Extinction.

After year 100, the directional trend, the increase in variability of climate, and the increased occurrence and severity of point extremes led to noticeable fluctuations in population size over time (Fig. 1). Twenty-one per cent of the 34 560 simulation replicates went extinct. As expected, risk of extinction increased with higher frequency and severity of point extreme events, older age at first reproduction, and fewer offspring produced per mating pair (Fig. 2). Given an expected yearly number of offspring per mating pair, the proportion of replicates that went extinct increased approximately linearly with age at first reproduction (Fig. 2a). When considering all replicates or only the most extreme scenario, for a fixed expected number of offspring produced per mating pair, increasing age at first reproduction by one year would increase the probability of going extinct by approximately 10% (Fig. 2a). The combination of relative high frequency and high severity of extreme events led to a noticeably higher proportion of replicates that went extinct (Fig. 2b). Among the replicates that went extinct, 16% of them were not affected by a point extreme in the 10 years before extinction and 34% in the 5 years preceding extinction.

**Figure 2.**
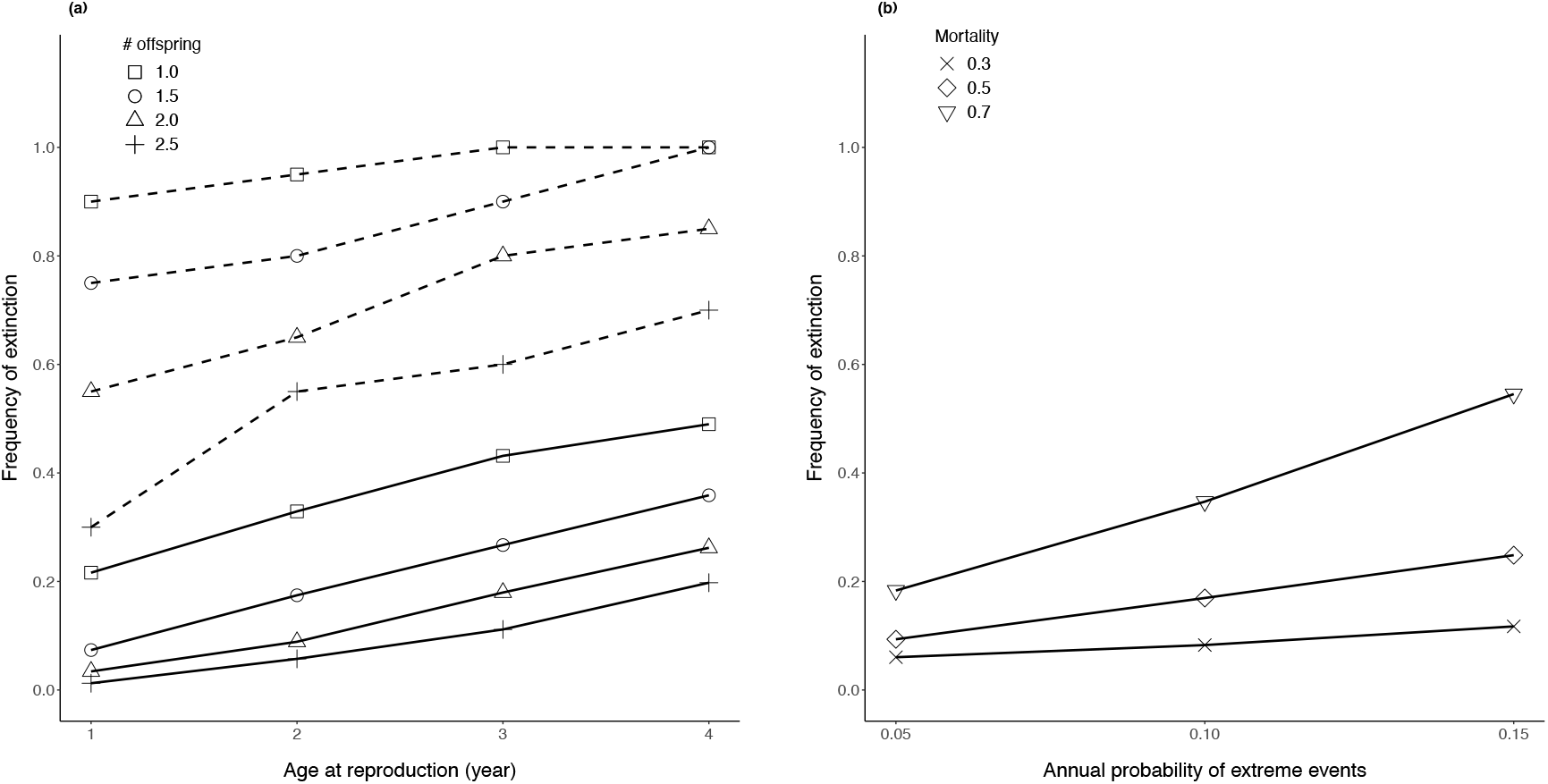
Extinction probability (number of populations going extinct divided by the number of replicates run for a given set of parameter values) for scenarios of: (a) different ages at first reproduction and expected yearly number of offspring per mating pair (solid line: all replicates; dashed line = most extreme environment, i.e., p(*E*_a_) = 15 10^-2^, *m*(*E*_a_) = 0.7, *β_μ,Θ_* = 5 10^-2^, *β_σ,Θ_* = 1.5 10^-2^); (b) different probability of point extremes and probability of dying from point extremes.

In populations that persisted until the end of simulation time, the GLM and GAM models that included *n_low* provided a good prediction of the total number of alleles at the end of simulation time (Table 2). The total number of years with population size smaller than 150 individuals had a strong, negative non-linear effect on the total number of alleles at the end of simulation time (Fig. 3). Models that did not include *n_low* had much lower predictive performance (Table 2).

**Table 2.** Performance of Ordinary Least-Squares regression models (OLS), Generalized Additive Models (GAM) and Random Forest models (RF) when predicting the total number of alleles at the end of simulation time for the replicates that did not go extinct. All populations started with 10 alleles for each of the 10 loci for a total of 100 unique alleles. Full models include as predictors *n_low, β_σ,Θ_, β_μ,Θ_*, p(*E*_a_), *m_E_, a*_f_, *λ*_o_, and *s. n_low* is the number of years the population was below 150 individuals and all other symbols are as in Table 1 (results were similar with *tr_low* = 100 or 50 and are reported in the computer code associated with this paper). The training data set had 21 523 replicates (80% of the replicates that survived up to the end of simulation time) and the test data set had 5 371 replicates. *R*^2^ was calculated with respect to the 1:1 predicted-observed line; *MAE* is the mean absolute error calculated over the whole test data set. Over the entire data set (training and test), the number of alleles at the end of simulation time was [mean ± sd] 79.84 ± 21.11.

| *Model* | $R^2$ | $MAE$ |
| --- | --- | --- |
| RF_*full* | 0.82 | 6.6 |
| RF_*red* | 0.24 | 15.0 |
| GAM_*full* | 0.78 | 7.8 |
| OLS_*full* | 0.71 | 8.8 |
| OLS_*red* | 0.22 | 15.2 |

**Figure 3.**
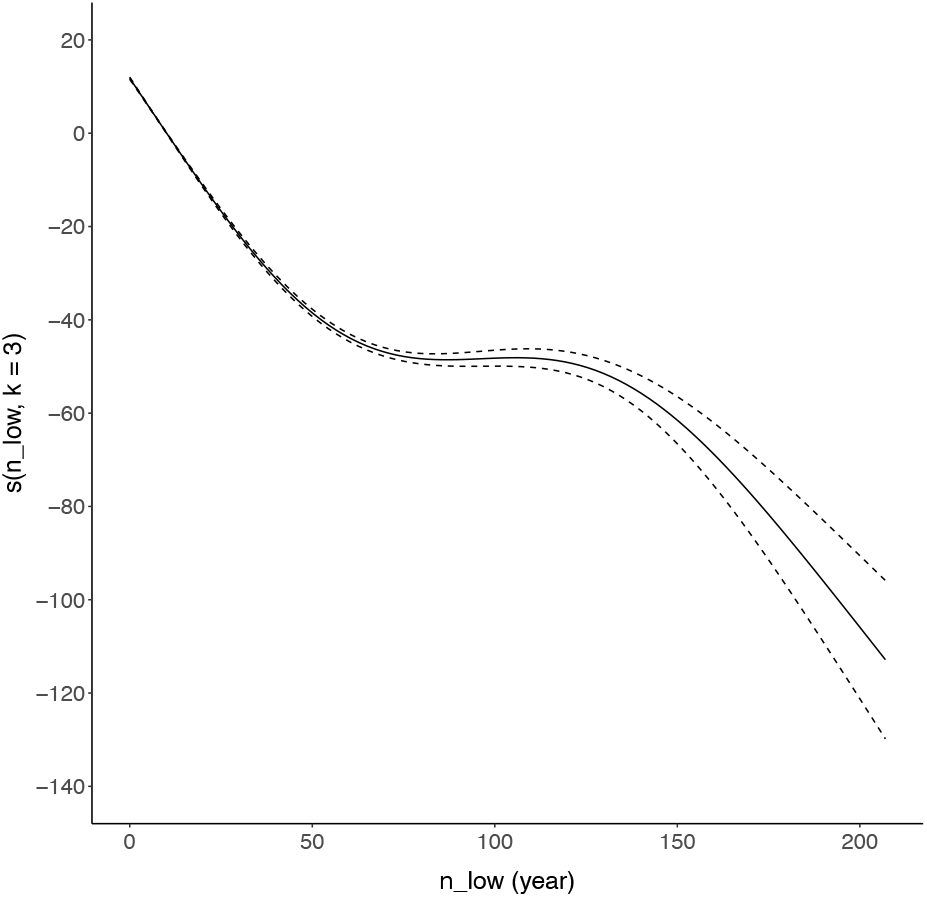
Partial non-linear effect of the total number of years in which the population is below 150 individuals during simulation time (*n_low*) on the total number of alleles at the end of simulation time, as found from the Generalized Additive Model of Table 2. The GAM algorithm found *k* = 3 as the optimal number of degrees of freedom for the spline (i.e., a cubic). Model details and results are in the in the computer code associated with this paper.

Models for differences in allelic frequency of more or less theoretically advantageous alleles were able to explain less than 5% of the variance of the target variable (Fig. 4). The average value of population-mean phenotype 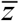 in the last ten years of simulation time was negatively correlated with the difference in frequency of top and bottom 10% of alleles according to their allelic value at the end of simulation time (*r* = 0.71, *p* < 0.01).

**Figure 4.**
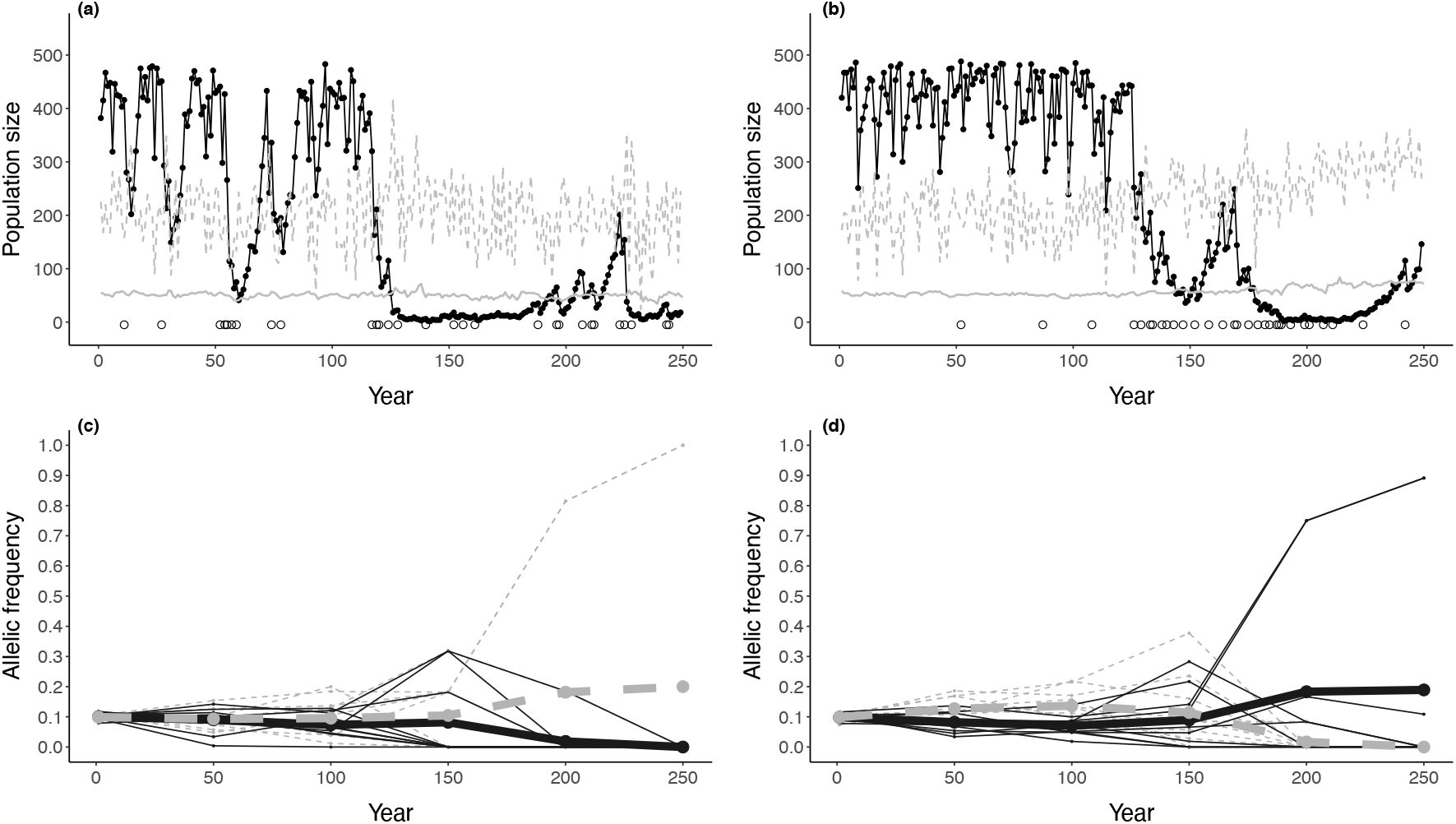
Panels (a-b): population dynamics of two simulation replicates (details about the replicates are in the computer code associated with this paper). Lines and symbols are as in Figure 1. Panels (c-d) dynamics of allelic frequency for the populations in panels (a-b) (a→c, b→d). Thin dashed gray lines represent the allelic frequencies measured every 50 years for alleles in the bottom 10% of allelic values (for (c): *n* = 10, allelic value [mean ± sd] = −0.27 ± 0.08; (d) *n* = 10, allelic value = −0.21 ± 0.04) and thin solid black lines represent the top 10% (for (c): *n* = 10, allelic value = 0.28 ± 0.08; (d) 0.25 ± 0.06). Thick dashed gray lines represent the average frequency in the population of alleles in the bottom 10% of allelic values and thick solid black lines represent the average frequency of the top 10%. In (c), the alleles in the bottom 10% of allelic values were more frequent in the population at the end of simulation time than alleles in the top 10%, and *vice versa* in panel (d).

The GLM, GAM, and RF models fitted on the training data sets had similar high predictive accuracy and low false positive and negative rates when predicting extinction or survival in the 10-year “prediction window” (Table 3). For more than 98% of replicates included in the test data set, the GLM, GAM, and RF models provided the same prediction of either extinction or persistence (Fig. 5). In the GAM model, only minimum population size in the observation window had a strong non-linear (negative) effect on the log-odds of population extinction. Likewise, minimum population size was the most important predictor in the RF model (Fig. 6a). The models without minimum and mean population size as predictors had fairly low accuracy (Table 3). The RF model without minimum and mean population size as predictors found age at reproduction and expected yearly number of offspring per mating pair as the most important predictors of contemporary extinction (Fig. 6b).

**Table 3.** Generalized Linear Model with logit link function for prediction of extinction (1)/persistence (0) of a population over a 10-year period (“prediction window”) with predictors minimum population size min(*N*), mean population size 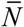, maximum value of the optimum phenotype max(*Θ*), maximum and minimum mean population-level maladaptation (or “extremeness” of the climate extreme) max(*Θ(t) − z(t))* and mean(*Θ(t)-z(t))*, and occurrence of a point extreme *E*, all measured in the 10 years before the start of the “prediction window” (i.e., in the “observation window”), along with selection strength *s*, age at first reproduction *a*_f_ and expected yearly number of offspring per pair *λ*_0_ (*full* model). In the *reduced* model, I excluded min(*N*) and 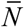. All predictors were standardized (*s* and *E* are categorical variables with two levels each and they were not standardized) and I report mean estimate and standard error of the regression coefficients for the GLM models (estimate, standard errors and significance where applicable for the GAM and RF models are in the computer code associated with this paper). The GLM, GAM, and RF models were trained on 39 547 replicates (36% went extinct) and tested on 4 600 replicates (50% went extinct). For the full and reduced GLM models, when using the optimal cutoffs (*full* = 0.54, *reduced* = 0.35), accuracy when tested on the validation data set was 97% for the *full* model and 77% for the *reduced* model, false positive rate 0.03 and 0.26, false negative rate 0.03 and 0.19. For the GAM models (details in computer code associated with this article), optimal cutoffs were 0.38 for the *full* model and 0.35 for the *reduced* model, accuracy 97% and 77%, false positive rate 0.03 and 0.25, false negative rate 0.03 and 0.20. For the RF models, accuracy when tested on the test data set was 97% for the *full* model and 72% for the *reduced* model, false positive rate 0.02 and 0.16, false negative rate 0.04 and 0.38.

|  | *full* | *reduced* |
| --- | --- | --- |
| *Intercept* | -5.63(0.14) | -2.39(0.03) |
| $\min(N)$ | -5.41(0.20) | - |
| $\bar{N}$ | -1.34(0.10) | - |
| $S$ | 0.62(0.06) | 1.02(0.03) |
| $\max(\Theta)$ | -0.14(0.04) | -0.11(0.02) |
| $\lambda_0$ | -0.42(0.03) | -0.89(0.01) |
| $a_f$ | 0.42(0.03) | 0.80(0.01) |
| $E$ | 0.64(0.07) | 1.50(0.03) |
| $\max(\Theta(t) - z(t))$ | 0.06(0.05) | 0.28(0.02) |
| $\text{mean}(\Theta(t) - z(t))$ | 0.13(0.04) | 0.62(0.03) |

**Figure 5.**
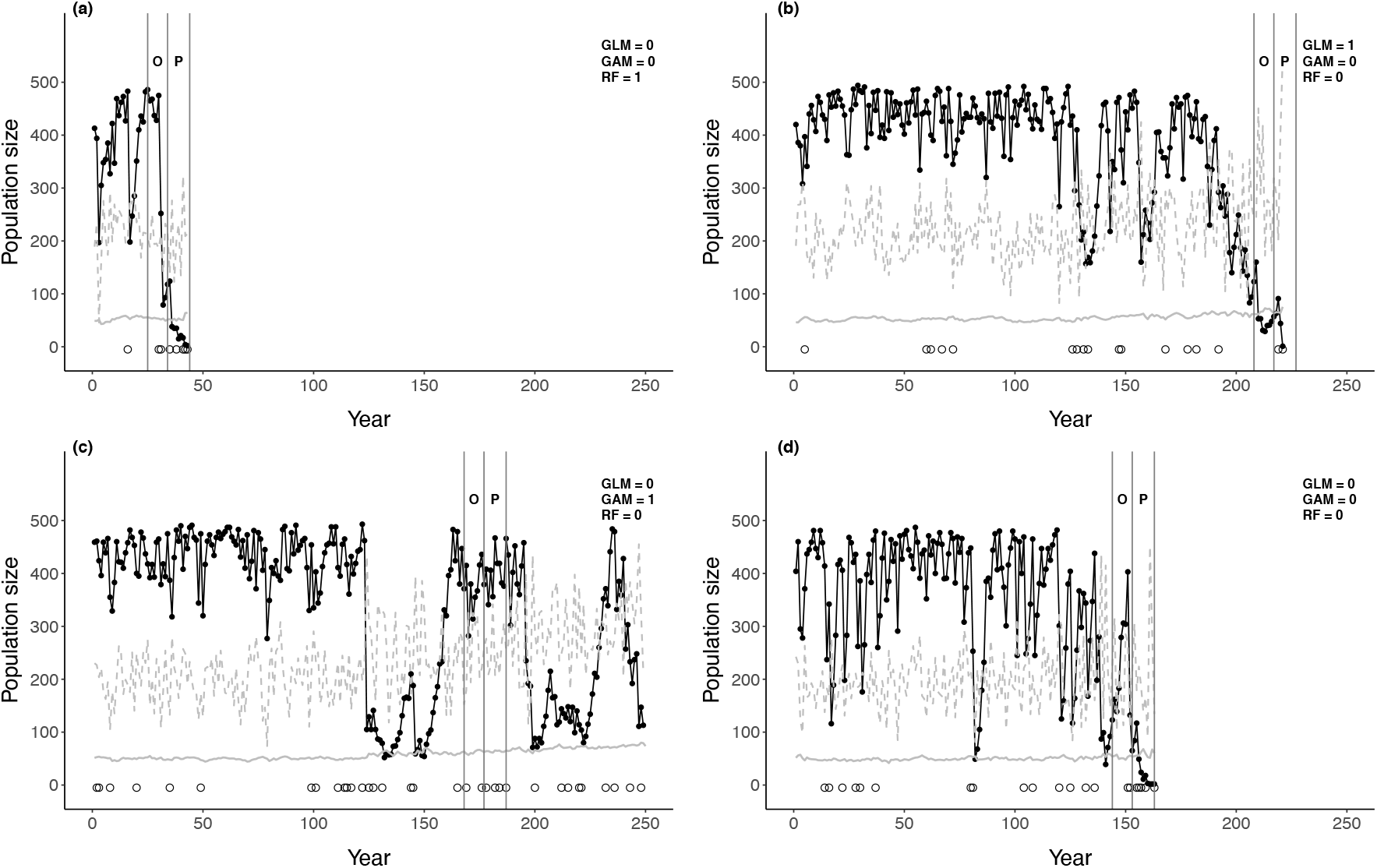
Examples of Generalized Linear Model (GLM), Generalized Additive Model (GAM), and Random Forest (RF) predictions of extinction in a “prediction window” based on climate and population traits observed in a “observation window”. Circles are point extremes, the black solid line and black points represent population size, the gray dashed line is the optimum phenotype re-scaled for graphical purposes. The letter O denotes the “observation window” and P the “prediction window”. For each model in the legend, 1 means the model predicted extinction (e.g., RF = 1 means that the RF model predicted extinction) and 0 otherwise; in panel (d), all models predicted extinction, but the replicate did not go extinct. In panel (a) only the RF model predicted extinction and the replicate went extinct, in (b) only the GLM model predicted extinction and the replicate went extinct, and in (c) only the GAM model predicted extinction and the replicate did not go extinct.

**Figure 6.**
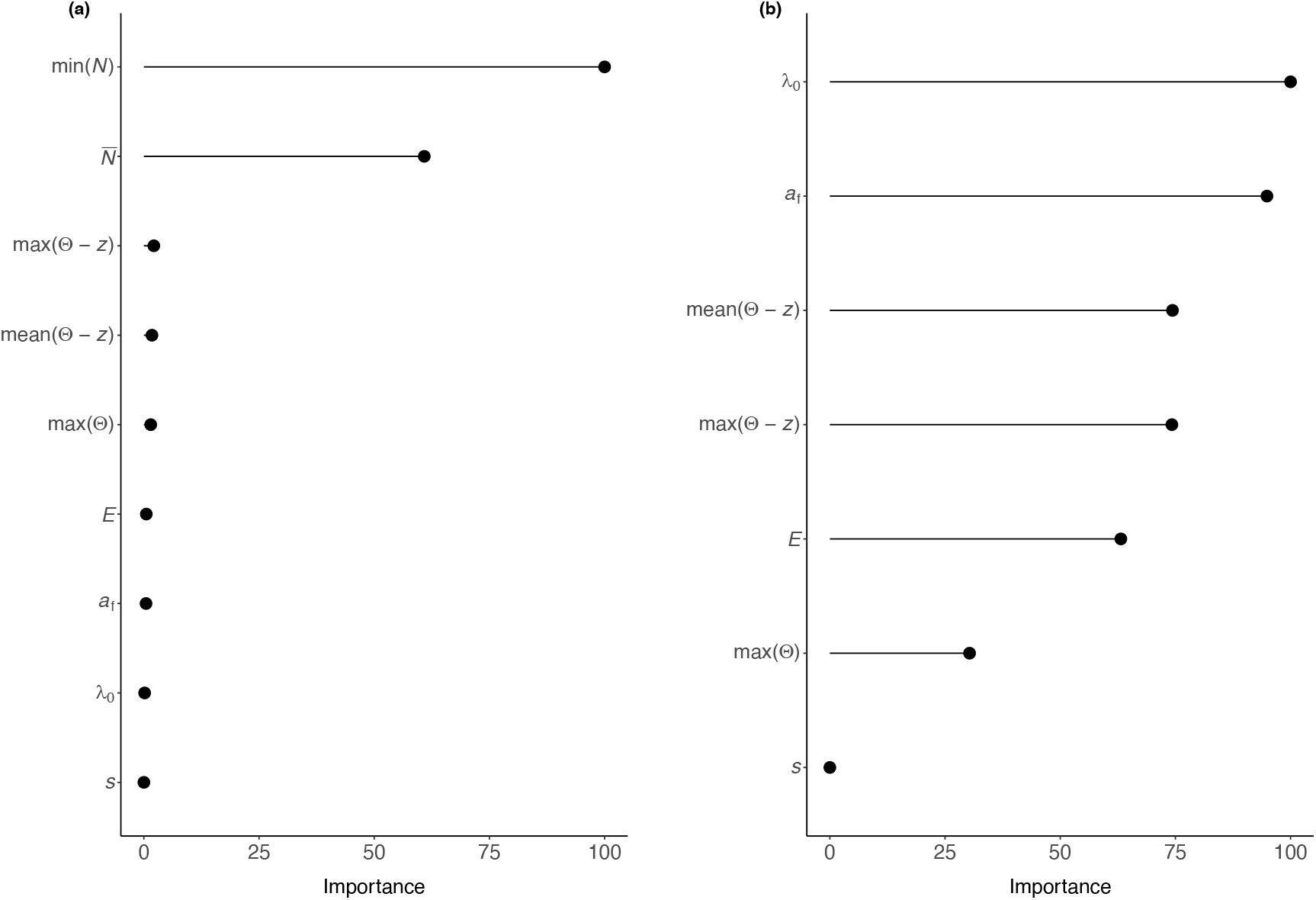
Variable importance for the full (panel (a)) and reduced (i.e., without min(*N*) and 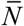, panel (b)) Random Forest models for prediction of extinction in the 10-year “prediction window” based on climate/environmental and population traits measured in the 10-year “observation window”. To compute variable importance, for each tree, the prediction accuracy on the out-of-bag portion of the data is recorded, and the same is done after permuting each predictor. The difference between the two accuracies are then averaged over all trees and normalized by the standard error. The most important predictor is assigned the value of 100 and the other predictors are scaled accordingly.

## 4 Discussion

Understanding and predicting the effects of extreme events on risk of extinction and population and genetic dynamics of natural populations is critical for both population forecasting and for managing human intervention in an increasingly more extreme world.

The results of the numerical simulations showed additive effects—with largely no interaction effects—between age at first reproduction and fecundity on risk of extinction for the range of values I simulated. In the replicates that survived up to the end of simulation time, the total number of years in which the population was at a small size was a good predictor of allelic richness for the quantitative trait under selection. The population frequency of theoretically advantageous alleles was strongly correlated with the mean value of the phenotype under selection but was otherwise largely unpredictable. Last, simple models including basic demographic and vital rates information, along with climate and environmental data, provided excellent predictions of contemporary risk of population extinction.

### 4.1 Life histories

Life-history theory predicts a prevalence of fast life histories in environments in which extreme events occur (Winemiller 2005). Fast life histories—usually defined as including a combination of faster body growth early in life, younger age at maturity, and higher reproductive effort early in the reproductive life of the individual—should allow for faster population growth rate after a drastic reduction in population size, an adaptive life-history strategy when the population is at risk of extinction.

For many animal species that frequently experience the often-catastrophic effects of extreme climate or point events—and have limited movement range for physiological and behavioral reasons or for the environment they inhabit (e.g., freshwater fish in mountain streams, insects)—offspring production is typically very high compared to the habitat carrying capacity. Thus, offspring production is not usually what is limiting population recovery, i.e. the re-establishment of the pre-event population size. However, when after a climate or point extreme event the population is reduced to such low numbers that the population is at immediate risk of extinction, younger age at first reproduction for the surviving individuals could be the life-history trait that makes the difference between population persistence or extinction. Vincenzi et al. (2017) found that fish born after flash floods had younger mean age at reproduction than fish born before flash floods; they hypothesized that younger age at reproduction after flash floods was due to a combination of faster growth due to lower population density and fewer older fish competing for mates. However, younger age at sexual maturity and higher energetic investment in offspring production often come at the cost of shorter life expectancy (Fay et al. 2016), and life histories that are adaptive after population crashes can show lower fitness in steady-state conditions (Vincenzi et al. 2012, 2014).

### 4.2 Prediction of genetic dynamics and of population extinction

Populations experiencing recurrent bottlenecks are expected to have their genetic pool eroded over time. The erosion of the genetic pool should be noticeable in particular in the loss of allelic richness, even after a single bottleneck event (Allendorf 1986). I found that the extent of population bottlenecks, which I operationalized as the number of years in which population size was relatively small with respect to the habitat carrying capacity and species numerical potential, was a strong predictor of allelic richness for the quantitative trait under selection in the simulations. These results appear to be partially consistent with recent empirical results. Vincenzi et al. (2017) found an increase in the proportion of fixed alleles in year-classes of two trout populations born after flash floods that caused massive mortalities. Poff et al. (2018) tested predictions about population genomic change in aquatic insects living in Colorado, US, mountain streams after a 1-in-500-year rainfall event. They found that allelic richness at presumably neutral loci declined after the event only in two out of six species analyzed. Moderate reduction of allelic diversity after strong bottlenecks might be attributable to the particular demographic history of the populations that are investigated; according to Bouzat (2010), one can expect that populations experiencing recurrent bottlenecks might have had their genetic pool already eroded over time, which would decrease the effectiveness of both purifying selection and random allele loss. In addition, the loss of alleles could be greater in species—like salmonids——with high variance in reproductive success among adults (i.e., greater than Poisson variance in reproductive success), although high variance in reproductive success has been found to bias (i.e., make false positives more likely) empirical investigations of genetic bottlenecks (Hoban et al. 2013). Moreover, in the simulations of population dynamics, we started from 10 unique alleles for each of the 10 loci, and the high initial allelic diversity may explain the strong relationship that was found between the temporal extent of bottlenecks and allelic richness.

On the other hand, a model including age at first reproduction and fecundity of the species, along with some traits of the environment, was not able to predict the dynamics of the population frequencies of more advantageous alleles for the quantitative trait under selection. However, a strong correlation was found between the frequencies of more or less advantageous alleles at the end of simulation time and the average value of the phenotype under selection. This result seems to at least suggest that, although the fate of alleles is difficult to predict from just the coarse-grain description of the environment and of the species, even in an extreme environment that also causes trait-independent mass mortalities, a shift in the phenotype is likely to be caused by the increased prevalence of the most advantageous alleles.

One of the foundational tenets of conservation biology is that small, fragmented populations should be considered locally vulnerable to extinction—even more critically so when affected by highly variable climatic conditions and other environmental disturbances. The conditions that led to the extinction of a population or species can almost always be understood retrospectively, but forecasting extinction, especially over contemporary time horizons, is much more challenging. In this work, I found that the most important predictors of contemporary extinctions were mean and minimum population size measured in the few years before the “prediction window”. However, it was not uncommon for the simulated populations to swiftly rebound after collapses in numbers, and age at reproduction and yearly fecundity were the most important predictors of extinction when measures of population size were not included in the model. This result highlights the importance of age at first reproduction and yearly fecundity for population persistence in highly stochastic environments, as also suggested by theoretical (Bürger and Lynch 1995) and experimental (Griffen and Drake 2008) studies. However, other genetic challenges not accounted for in my simulation model are likely to be encountered by populations that decline to very small numbers, such as a reduction of viability and/or fecundity due to either inbreeding or the expression of deleterious alleles (Willi et al. 2006).

Although it may appear from the simulation results that vital rates and environmental conditions play a small role in models that predict contemporary risk of extinction, those rates and conditions heavily contribute to determining population size in the “observation window”; for example, populations with higher fecundity and younger age at first reproduction are less likely to remain at low population sizes than populations with lower fecundity and delayed sexual maturity. Likewise, a more extreme environment (e.g., greater selection strength and more frequent and/or severe climate and point extremes) tends to decrease average population size, either due to acute events that kill individuals or constant recovery from population crashes.

Luck also plays a major role in determining whether a population will recover after a population crash. Vincenzi et al. (2017) found that the almost-complete recovery of a trout population that was reduced to a handful of individuals after a flash flood was due to the large production of offspring by a single mating pair. Considering the high variance in adult reproductive success in trout populations—which is at least partially due to differences in individual “quality” (Auld et al. 2019)—had the mating pair been killed or displaced during the flash flood, population recovery would have suddenly become much less likely.

### 4.3 Modeling considerations

Modeling and simulation approaches can help understand the effects of multiple extreme stressors on the contemporary risk of extinction of species and can be used to guide or support both the set-up of ecologically relevant experimental designs and the interpretation of biological responses to multiple stressors.

However, models of population and genetic dynamics are limited in their scope of prediction by both the understanding of the biology and ecology of the species and the availability of data to parametrize, train, and test models, along with modeling choices (e.g., explicit or implicit modeling of alleles, clustered or “well-mixed” individuals or populations). Although often intuitive, it is nevertheless important to remind ourselves that the simulation results of modeling exercises depend on, first, our biological and ecological understanding, and, second, the simplified modeling of the species and the environment they inhabit. For instance, general trade-offs between allocation of resources to competing physiological functions are not only often intrinsically challenging to model, but they may also vary over time and space for the same species or population. That is, even when there is a qualitative understanding of biological or ecological processes, the parameterization and choice of parameter values for the model may be too uncertain to provide actionable predictions.

Trade-offs between model accuracy and interpretability also need to be taken into account when developing models of population and genetic dynamics. Accuracy describes the ability of a model to explain observed data and make correct predictions, while interpretability concerns to what degree the model allows for understanding processes. Often, there is a trade-off between accuracy and interpretability: more complex models are usually opaque, while more interpretable models often do not provide the same accuracy or predictive power of more complex models (Breiman 2001b). Then, although intuitively more complex models are expected to provide more accurate predictions of risk of extinction or population and genetic dynamics, this is not always the case. For instance, Ward et al. (2014) tested the predictive performance of short-term forecasting models of population abundance of varying complexity. They found that more complex models often performed worse than simpler models, which simply treated the most recent observation as the forecast. In their case, the estimation of even a small number of parameters imposed a high cost while providing little benefit for short-term forecasting of population abundance. However, when there was a clear signal of cyclic dynamics, more complex models were able to more accurately predict future population sizes.

As always, the purpose of a scientific investigation should drive model formulation, the type and amount of data collected, and the acceptable uncertainty of model predictions.

## Ethics

Not applicable

## Data accessibility

Data and relevant code for this research work are stored in GitHub: https://github.com/simonevincenzi/Contemporary_Extinction

## Authors’ Contributions

Simone Vincenzi conceived the ideas, designed the methods, run the analyses, and wrote the manuscript.

## Acknowledgements

Simone Vincenzi developed the idea behind this work while walking the Malecón of La Habana, Cuba, and finished writing the manuscript after the 2020 California fires that imperiled the town where he resides (Santa Cruz, CA, US) and the COVID-19 pandemic highlighted the importance of proper simulation modeling for basic science and policy, and the increasing threat of extreme events for people and nature.

